# Preconceptional social isolation alters inflammatory and transcriptional signalling across the placenta–fetal brain axis

**DOI:** 10.64898/2026.10.01.755920

**Authors:** Monica Iachizzi, José Ruiz Rodriguez, Felisa Herrero, Ron Schär, Annamaria Cattaneo, Juliet Richetto

**Affiliations:** Institute of Pharmacology and Toxicology, University of Zurich-Vetsuisse, Zurich, Switzerland; Neuroscience Center Zurich, University of Zurich and ETH Zurich, Zurich, Switzerland; Functional Genomics Center Zurich, University of Zurich and ETH Zurich, Zurich, Switzerland; Department of Pharmacological and Biomolecular Sciences, University of Milan, Milan, Italy; Biological Psychiatry Laboratory, IRCCS Istituto Centro San Giovanni di Dio Fatebenefratelli, Brescia, Italy

## Abstract

Maternal depression and psychosocial adversity during pregnancy are associated with altered offspring neurodevelopment, yet the molecular processes linking maternal state to the developing fetus remain incompletely understood. The placenta is positioned to sense and respond to maternal physiological signals, but how placental molecular responses relate to those in the developing brain is unclear. Here, we used a mouse model of prolonged social isolation initiated before conception and maintained throughout pregnancy to investigate inflammatory and transcriptional alterations across matched placenta and fetal-brain samples at gestational day 18. Maternal social isolation altered cytokine profiles in both tissues and induced extensive transcriptional changes in the placenta, with a more restricted response in the fetal brain. Functional analyses identified alterations in placental immune, metabolic and transport-related programmes and fetal-brain immune, neuronal and proliferative processes. Although sex-stratified analyses revealed different transcriptional profiles in male and female tissues, formal maternal housing × fetal-sex interaction testing did not identify significant gene-level interaction effects. Cross-tissue co-expression analysis initially identified extensive covariation between social-isolation-responsive placental and fetal-brain modules; however, most associations were attenuated after accounting for their shared relationship with maternal housing. An exposure-independent network analysis nevertheless identified an inverse pregnancy-level association between a placental module enriched for immune and interferon-related signalling and a fetal-brain module enriched for cell-cycle and proliferative processes. Together, these findings show that prolonged maternal social adversity is associated with coordinated but tissue-distinct molecular alterations across the placenta–fetal brain axis and demonstrate the importance of distinguishing parallel responses to a common maternal environment from direct cross-tissue molecular coupling.

## Introduction

Maternal depression during pregnancy is one of the most common early-life adversities, yet the biological pathways through which it shapes fetal development remain poorly defined. Clinically significant depressive symptoms affect approximately one in five pregnancies, while major depressive disorder affects a substantial subset of pregnant women^1–4^. Beyond maternal suffering, antenatal depression is associated with adverse birth outcomes and with increased offspring vulnerability to emotional, cognitive and behavioural difficulties and later psychopathology^5–7^. Antenatal depression is also accompanied, in at least a subset of pregnancies, by low-grade immune activation, including alterations in interleukin-6 (IL-6), tumour necrosis factor-α (TNFα) and C-reactive protein, together with dysregulation of hypothalamic–pituitary–adrenal (HPA)-axis and glucocorticoid signalling^8–12^. The fetus may therefore develop in the context of coordinated changes in maternal inflammatory, endocrine and metabolic physiology^10–17^. How these maternal alterations become biologically embedded during fetal development remains unresolved.

The placenta is the principal interface through which maternal physiological signals are sensed, filtered and translated into the fetal environment. It coordinates nutrient and xenobiotic transport, vascular development, and endocrine and immune communication, thereby regulating fetal exposure to maternal hormones, nutrients and inflammatory mediators^18–21^. Rather than acting as a passive barrier, the placenta can buffer, amplify or transform maternal physiological signals. Placental immune-sensing pathways respond to maternal inflammatory states and can engage cytokine and NF-κB-dependent signalling in trophoblast cells^22,23^. Such responses may alter trophoblast function, angiogenic signalling, nutrient transport and placental endocrine and immune communication, providing potential routes through which depression-associated maternal physiology could influence fetal development^24,25^.

Human placental studies have associated maternal stress or depressive symptoms with transcriptional alterations involving immune regulation, cellular stress and protein synthesis, metabolism, trophoblast differentiation, vascular function and nutrient transport^26–31^. These findings support the placenta as a potential site at which maternal psychological and physiological states become biologically embedded during pregnancy. However, human studies remain inherently limited by heterogeneity in diagnosis and symptom timing, medication exposure, comorbidities, delivery and tissue sampling, as well as placental cellular composition and other environmental and clinical factors. Experimental models are therefore required to isolate maternal exposures and examine molecular responses simultaneously across placenta and developing brain.

Experimental models have demonstrated that prenatal stress and maternal immune activation can alter offspring stress-axis regulation, neuroinflammatory signalling, neurogenesis and later affective behaviour ^32–36^. However, many models employ physical or acute stressors during restricted gestational windows, whereas social adversity is an important contributor to depression risk in humans and vulnerability frequently predates pregnancy^37^. Preconceptional stress may therefore represent an important but comparatively understudied component of maternal vulnerability. Notably, experimental preconception-stress studies have identified transcriptional alterations in both placenta and fetal brain and have reported different patterns of response in male and female tissues^38^. Critically, however, relatively few studies have examined molecular responses simultaneously across matched placenta and fetal brain, and it remains unclear to what extent transcriptional programmes in the two tissues covary within pregnancy.

Fetal sex nevertheless represents an important biological variable in placental and fetal responses to maternal adversity. Male and female placentas differ in baseline transcriptional, endocrine and immune characteristics^39,40^, and sex-stratified studies have reported divergent molecular responses to prenatal and preconceptional stress^38,41^. These observations highlight the importance of considering fetal sex when characterizing molecular responses across the placenta–brain axis, while distinguishing descriptive differences between sex-stratified responses from formal evidence of a sex-dependent exposure effect.

Here, we used a social isolation rearing (SIR) model to investigate how preconceptional social adversity that persists throughout pregnancy alters molecular signalling across the placenta–fetal brain axis. The model induces affective and physiological alterations in females through prolonged social isolation initiated before conception, and previous studies have demonstrated lasting, partly sex-dependent effects on offspring development^42,43^. We quantified inflammatory mediators and performed RNA sequencing in matched placenta and fetal-brain samples from male and female fetuses at late gestation. We combined differential-expression and functional-enrichment analyses with cross-tissue co-expression network analysis to determine whether maternal SIR induces coordinated molecular alterations across the two tissues. We further formally tested whether transcriptional responses to SIR differed by fetal sex. This design allowed us to characterize inflammatory and transcriptional responses to maternal SIR across the placenta–fetal brain axis and to identify candidate cross-tissue programmes associated with the maternal environment.

## Materials and Methods

### Animals

All animal procedures were conducted in accordance with Swiss legislation and were approved by the Cantonal Veterinary Authority of Zurich (animal licence ZH142/2024). C57BL/6N mice were maintained in individually ventilated cages under specific-pathogen-free conditions in a temperature- and humidity-controlled facility (21 ± 3 °C; 50 ± 10% relative humidity) under a reversed 12-h light–dark cycle (lights off 09:00–21:00). Food and water were available ad libitum. These housing conditions follow those established previously for the social isolation rearing model.

### Social isolation rearing and mating

Female C57BL/6N mice were randomly allocated at postnatal day (PND) 21 to either group housing (GRP; 3-4 animals per cage) or social isolation rearing (SIR; one animal per cage). Animals were maintained under these conditions for 10 weeks before mating. The SIR paradigm and mating procedure were based on our previously established model of preconceptional social adversity^42^.

Following the 10-week preconceptional housing period, females were subjected to timed mating with adult C57BL/6N males. Successful mating was determined by the presence of a vaginal plug and designated gestational day (GD) 0. Pregnant SIR females remained individually housed throughout gestation, whereas GRP females were maintained under group-housing conditions.

For the present study, seven pregnant dams per housing condition were included in the molecular analyses. One male and one female fetus from each pregnancy were selected for each tissue analysis, resulting in 28 placental and 28 fetal-brain samples (GRP: 7 male and 7 female; SIR: 7 male and 7 female).

### Tissue collection

At gestational day 18 (GD18), pregnant dams were anaesthetized with isoflurane and euthanized by decapitation. The uterus was rapidly removed and transferred to ice-cold phosphate-buffered saline (PBS). Individual conceptuses were dissected sequentially, with decidual tissue and yolk sac removed and the placenta separated from the fetus. Placentae were gently washed in ice-cold PBS to remove residual blood. Fetal brains were rapidly dissected on a surface cooled with dry ice. Placenta and corresponding fetal-brain samples were snap-frozen in liquid nitrogen and stored at −80 °C until processing. Fetal sex was determined by DNA-based genotyping to enable selection of one male and one female fetus, together with the corresponding placenta, from each dam. Genomic DNA was isolated from GD18 fetal tissue using an alkaline lysis protocol. Briefly, tissue was incubated in 75 µl lysis buffer (25 mM NaOH, 0.2 mM EDTA; pH 12) for 20 min at 95 °C and neutralized with 75 µl Tris-HCl (40 mM, pH 5). Sex was determined by real-time PCR using primers targeting the sex-determining region of the Y chromosome (SX), with GAPDH as an internal control. Amplification was performed using SsoAdvanced™ Universal SYBR® Green Supermix (Bio-Rad) on a CFX384 real-time PCR system (Bio-Rad), and amplification specificity was verified by melting-curve analysis. One male and one female fetus from each dam were selected for molecular analyses, with placental and fetal-brain samples retained as matched pairs from the same fetus.

### Cytokine measurements

Placental and fetal-brain tissues were thawed on ice in ice-cold complete lysis buffer (cOmplete™ Lysis-M, Sigma-Aldrich, Switzerland) and homogenized using a TissueLyser II (Qiagen) for 3 min at 20 Hz. Homogenates were centrifuged at 12,000 rpm for 20 min at 4 °C, and supernatants were collected and stored at −80 °C until analysis. Concentrations of interferon-γ (IFNγ), tumour necrosis factor-α (TNFα), interleukin-6 (IL-6) and interleukin-1β (IL-1β) were quantified using a Meso Scale Discovery (MSD) V-PLEX electrochemiluminescence multiplex assay for mouse cytokines (MSD, Rockville, MD, USA), according to the manufacturer’s instructions and established protocols^44,45^. Samples were analysed in duplicate and cytokine concentrations were calculated from plate-specific standard curves using MSD Discovery Workbench software, adjusted for sample dilution, and normalized to total protein content. For each tissue, cytokine concentrations were analysed using two-way ANOVA with maternal housing (GRP/SIR) and fetal sex (male/female) as factors, with statistical significance set at *p* < 0.05.

### RNA extraction and RNA sequencing

Placental and fetal-brain tissues were thawed on ice in ice-cold RLT Buffer (Qiagen, Switzerland) and homogenized using a TissueLyser II (Qiagen) for 3 min at 20 Hz. Homogenates were centrifuged at 12,000 rpm for 20 min at 4 °C, and total DNA and RNA were co-extracted using the AllPrep DNA/RNA Mini Kit (Qiagen, Switzerland). RNA and DNA concentration and integrity were assessed using an Agilent Bioanalyzer, and samples with RNA integrity number (RIN) and DNA integrity number (DIN) > 7 were retained for downstream analyses.

RNA-sequencing libraries were prepared from 500 ng total RNA using the Illumina Stranded mRNA library preparation kit (Illumina). Briefly, poly(A)-enriched RNA was fragmented, reverse-transcribed into double-stranded cDNA and ligated to sequencing adapters, followed by PCR enrichment. Library quality and fragment-size distribution were assessed using an Agilent 2100 Bioanalyzer. Libraries were pooled and sequenced on an Illumina NovaSeq X Plus to an average depth of approximately 20 million 150-bp paired-end reads per sample. Raw sequencing reads were subjected to quality control and transcript abundance was quantified using the SUSHI data-analysis framework at the Functional Genomics Center Zurich. Transcript quantification was performed using Kallisto against the mouse reference transcriptome based on GENCODE release M31, and analyses were restricted to protein-coding genes.

Differential gene-expression analyses were initially performed separately for male and female placental and fetal-brain samples, comparing SIR with GRP offspring within each sex. Differentially expressed genes (DEGs) were defined using a false-discovery-rate (FDR)-adjusted p value < 0.05. The sex-resolved analyses comprised seven independent pregnancies per housing condition and fetal sex.

### Maternal housing × fetal-sex interaction analysis

To formally determine whether the transcriptional response to maternal SIR differed according to fetal sex, an interaction analysis was performed separately in placenta and fetal brain using all 28 samples per tissue. Because each dam contributed one male and one female fetus, the statistical model accounted for the paired litter structure.

Differential expression was analysed using DESeq2 with dam modelled as a fixed effect nested within maternal housing, using the design:

∼ housing + housing:dam + housing:sex

The maternal housing × fetal-sex interaction therefore tested whether the transcriptional response to SIR differed between male and female fetuses, corresponding to (SIRmale − GRPmale) − (SIRfemale − GRPfemale). As an independent analytical approach, interaction effects were additionally evaluated using limma-voom with dam included as a blocking factor. Multiple-testing correction was performed using the Benjamini–Hochberg procedure.

### Pathway and functional enrichment analyses

Canonical pathway enrichment and predicted pathway activity were assessed using Ingenuity Pathway Analysis (IPA; QIAGEN, Redwood City, CA, USA). Analyses were performed separately for placental and fetal-brain DEGs from male and female fetuses. Enrichment significance was determined using a right-tailed Fisher’s exact test, and IPA activation z scores were used, where available, to infer predicted pathway activation or inhibition. IPA Upstream Regulator Analysis was additionally used to identify candidate regulators whose known downstream targets showed expression patterns consistent with predicted regulator activation or inhibition. Upstream-regulator predictions were evaluated based on enrichment significance and IPA activation z scores.

Gene Ontology Biological Process (GO-BP) over-representation analysis was additionally performed separately for upregulated and downregulated DEGs from each tissue and sex. Enrichment was calculated relative to all expressed genes, and multiple-testing correction was performed using the Benjamini–Hochberg method. Terms with adjusted p < 0.05 were considered significantly enriched.

### Weighted gene co-expression network analysis

Weighted gene co-expression network analysis (WGCNA) was used to investigate coordinated transcriptional programmes across matched placenta and fetal-brain samples. Two complementary approaches were performed.

First, an exploratory DEG-selected analysis was performed using genes identified as differentially expressed in the sex-stratified SIR versus GRP contrasts. Separate networks were constructed using all DEGs and the upregulated and downregulated subsets, and analyses were performed across all samples and separately by fetal sex. Each co-expression module was summarized by its first principal component, termed the module eigengene (ME), and placental and fetal-brain MEs were correlated across matched samples. Because the genes used to construct these networks had themselves been selected on the basis of their response to SIR, significant cross-tissue associations were subsequently reassessed after adjustment for housing condition to determine the extent to which correlations reflected the common transcriptional response to maternal housing rather than exposure-independent cross-tissue covariation.

Second, an exposure-independent WGCNA was performed to identify cross-tissue associations without selecting genes according to their response to SIR. For each tissue, the 5,000 most variably expressed genes were selected independently of housing condition and used for network construction. Soft-thresholding powers were selected automatically during network construction; a power of 6 was selected for 8 of the 12 networks. As a sensitivity analysis, networks were additionally reconstructed using a soft-thresholding power of 8, as recommended by WGCNA guidance for the corresponding sample size. These sensitivity analyses are summarized in Figure 4B.Genes with correlated expression profiles were grouped into modules and summarized by their MEs. Placental and fetal-brain MEs were then correlated across matched samples.

To distinguish cross-tissue covariation from correlations arising from common effects of maternal housing, partial correlations were calculated after adjustment for housing condition. Because male and female fetuses originating from the same dam were not statistically independent, statistical significance was additionally assessed using a litter-aware permutation procedure in which whole litters were permuted within housing condition over 10,000 iterations. P values from module-pair comparisons were corrected using the Benjamini–Hochberg procedure. Given the exploratory nature of the cross-tissue network analysis and the limited number of independent pregnancies, associations with FDR < 0.10 were retained for follow-up, with both conventional and litter-aware permutation-based FDR values reported.

For the exposure-independent module pair that remained associated after adjustment for housing and litter-aware permutation testing, the contribution of between- and within-litter variation was further examined by comparing correlations between litter-averaged module eigengenes with correlations between within-litter male–female ME differences. Genes belonging to modules showing evidence of cross-tissue association were subjected to GO Biological Process enrichment analysis to characterize the biological processes represented by each co-expression programme.

### Statistical analysis

Statistical analyses of cytokine concentrations were performed using GraphPad Prism/version 11. Cytokine data were analysed using two-way ANOVA with maternal housing and fetal sex as factors. RNA-sequencing analyses were corrected for multiple comparisons using the Benjamini–Hochberg FDR procedure as described above. Differentially expressed genes were defined at FDR < 0.05. Housing × fetal-sex interaction effects were additionally examined at FDR < 0.10 given the lower statistical power of interaction testing. For exploratory cross-tissue WGCNA, associations with FDR < 0.10 were retained for follow-up, with both conventional and litter-aware permutation-derived significance estimates reported where applicable. All statistical tests were two-sided unless otherwise specified.

## Results

### Maternal social isolation induces distinct inflammatory profiles in the placenta and fetal brain

We first quantified inflammatory mediators in matched placental and fetal brain tissues to determine whether SIR altered the inflammatory environment at the maternal–fetal interface and in the developing brain. As described in Figure 1A, female mice were housed either in groups (GRP) or in social isolation from postnatal day 21 and maintained under these conditions for 10 weeks before mating and throughout pregnancy. Placentae and fetal brains were collected from male and female fetuses at gestational day 18 (GD18) for cytokine and transcriptomic analyses.

**Figure 1.**
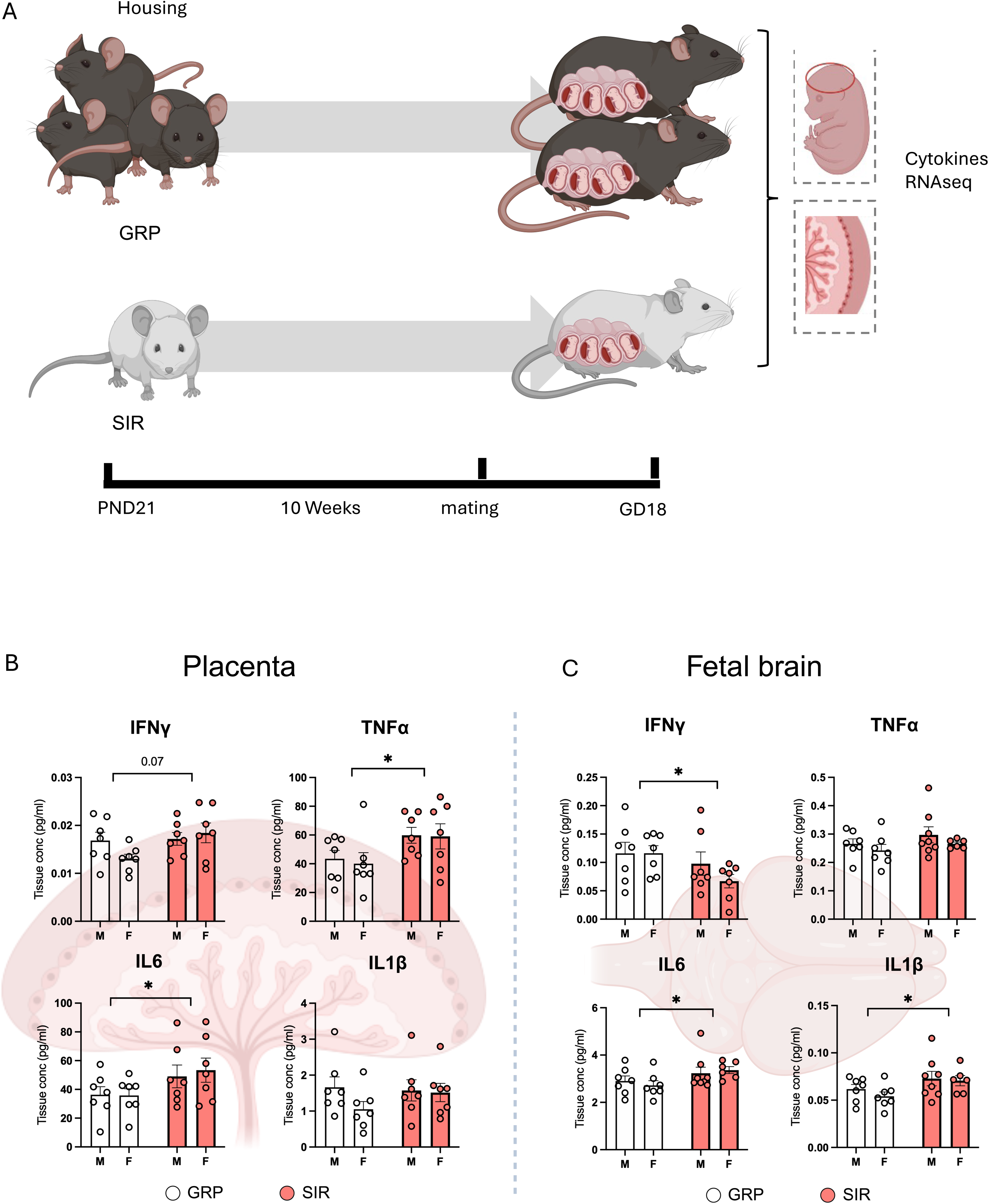
Maternal social isolation alters cytokine concentrations in the placenta and fetal brain. **A**) Experimental design. Female mice were housed either in groups (GRP) or individually under social isolation rearing (SIR) conditions from postnatal day 21. After 10 weeks, females were mated and maintained under their respective housing conditions throughout pregnancy. Placentae and fetal brains were collected from male and female fetuses at gestational day 18 (GD18) for cytokine measurements and RNA sequencing. **B**), Concentrations of interferon-γ (IFNγ), tumour necrosis factor-α (TNFα), interleukin-6 (IL-6) and interleukin-1β (IL-1β) in placental and fetal-brain tissue. White bars indicate GRP pregnancies and red bars indicate SIR pregnancies; M and F denote male and female fetuses, respectively. Individual points represent tissue samples from independent pregnancies, with one fetus of each sex analysed per dam (n = 7 pregnancies per housing condition and fetal sex). Data are presented as mean ± SEM. Cytokine concentrations were analysed using two-way ANOVA with maternal housing and fetal sex as factors. No significant main effects of fetal sex or maternal housing × fetal sex interactions were detected. Significant main effects of maternal housing were observed for placental TNFα (*p* = 0.019) and IL-6 (*p* = 0.036), and for fetal-brain IFNγ (*p* = 0.049), IL-6 (*p* = 0.035) and IL-1β (*p* = 0.030). Placental IFNγ showed a nonsignificant tendency to increase following SIR (*p* = 0.070). \**p* < 0.05; \*\**p* < 0.01; two-sided tests.

Maternal SIR altered cytokine concentrations in both tissues, although the affected cytokines and direction of change differed between the placenta and fetal brain. In the placenta, SIR increased tumour necrosis factor-α (TNFα; main effect of maternal housing, *F*_(1,24)_ = 6.28, *p* = 0.019; Figure 1B) and interleukin-6 (IL-6; main effect of maternal housing, *F*_(1,24)_ = 4.93, *p* = 0.036 Figure 1B), while interferon-γ tended to increase (IFNγ; main effect of maternal housing, *F*_(1,24)_ = 3.34, *p* = 0.07; Figure 1B). By contrast, placental IL-1β concentrations were not significantly altered by maternal housing (Figure 1B).

In the fetal brain, SIR was associated with lower IFNγ concentrations (main effect of maternal housing, *F*_(1,24)_ = 4.52, *p* = 0.049; Figure 1C), but higher IL-6 (main effect of maternal housing, *F*_(1,24)_ = 4.91, *p* = 0.035;Figure 1C) and IL-1β concentrations (main effect of maternal housing, *F*_(1,24)_ = 5.27, *p* = 0.030; Figure 1C). Fetal-brain TNFα concentrations did not differ significantly between maternal housing conditions (Figure 1C). No significant main effects of fetal sex or maternal housing × sex interactions were detected for any of the cytokines analysed. Together, these findings show that maternal SIR alters the inflammatory environments of both the placenta and fetal brain, with tissue-dependent effects on individual cytokines.

### Maternal SIR produces distinct transcriptional responses across placenta and fetal brain

We next performed RNA sequencing of matched placental and fetal-brain tissues to characterize the transcriptional response to maternal SIR. Initial analyses were stratified by fetal sex to characterize the transcriptional response within male and female tissues. In parallel, we formally tested maternal housing × fetal-sex interactions using a paired model accounting for the contribution of male and female fetuses from the same pregnancy. No individual transcript showed a significant housing × fetal-sex interaction after correction for multiple testing in either placenta or fetal brain (FDR < 0.10). Complete housing × fetal-sex interaction results are provided in Supplementary Data 2. We therefore interpret the sex-stratified analyses as describing the transcriptional response detected within each sex, rather than as evidence of statistically significant sex-specific effects. Maternal SIR was associated with extensive transcriptional alterations in the placenta. In female placentae, 1,270 differentially expressed genes (DEGs) were identified, of which 594 were upregulated and 676 were downregulated relative to GRP controls (Figure 2A). In male placentae, 1,124 DEGs were detected, comprising 605 upregulated and 519 downregulated genes (Figure 2C). By comparison, the fetal-brain transcriptional response was more restricted. In females, 129 DEGs were identified, including 54 upregulated and 75 downregulated genes (Figure 2B), whereas the male fetal brain contained 113 DEGs, of which 65 were upregulated and 48 were downregulated (Figure 2D). Complete differential-expression results for all four tissue–sex contrasts are provided in Supplementary Data 1.

**Figure 2.**
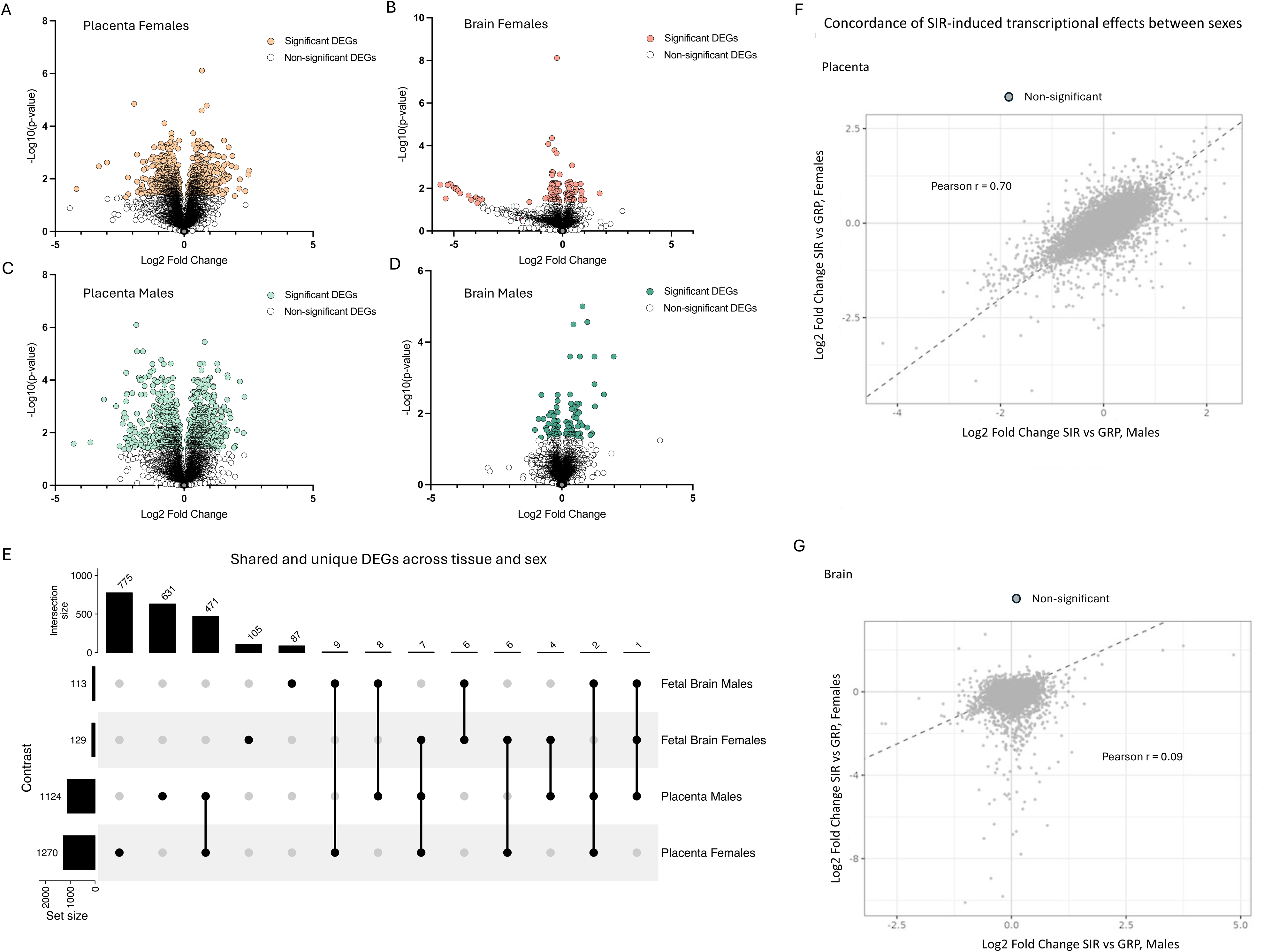
Maternal social isolation produces distinct transcriptional responses across the placenta and fetal brain. **A–D**, Volcano plots showing differential gene expression between SIR and GRP samples in female placentae (**A**), female fetal brains (**B**), male placentae (**C**) and male fetal brains (**D**). Coloured points indicate differentially expressed genes (DEGs) meeting the predefined significance criteria of *FDR-corrected p* < 0.05 threshold. n = 7 independent pregnancies per housing condition in each contrast. **E**, UpSet plot showing shared and unique DEGs across the four tissue–sex contrasts, irrespective of the direction of differential expression. Horizontal bars indicate the total number of DEGs in each contrast; vertical bars indicate the number of genes unique to, or shared among, the contrasts denoted by the connected points below. No DEG was common to all four contrasts. **F–G**, Concordance of SIR-associated gene-level effect sizes between male and female analyses in placenta (F) and fetal brain (G). Each point represents a gene, with log2 fold changes from the female SIR versus GRP contrast plotted against the corresponding log2 fold changes from the male contrast. Pearson correlation coefficients are shown. These analyses provide a descriptive assessment of effect-size concordance across sexes; formal housing × fetal-sex interactions were tested separately as described in the Methods. GRP, group housed; SIR, social isolation rearing; DEG, differentially expressed gene.

Intersection analysis revealed that most DEGs were restricted to an individual tissue–sex contrast (Figure 2E). Of these, 775 were detected exclusively in female placentae, 631 in male placentae, 105 in female fetal brains and 87 in male fetal brains. The greatest overlap was observed between the male and female placental responses, with 471 DEGs shared between the two placental contrasts. In comparison, only seven DEGs were shared between the male and female fetal-brain responses, six of which were restricted to the two fetal-brain contrasts and one of which was also differentially expressed in the male placenta. To further examine the similarity of the transcriptional response between male and female fetuses, we compared SIR-associated gene-level effect sizes between the two sex-stratified analyses (Figure 2F,G). In the placenta, SIR-associated log2 fold changes were positively correlated between males and females (r = 0.70), indicating substantial concordance in the direction and magnitude of placental transcriptional responses across sexes (Figure 2F). In contrast, effect-size concordance was weak in the fetal brain (r = 0.09; Figure 2G). These correlations provide a descriptive measure of similarity between the sex-stratified transcriptional responses and do not constitute evidence of a sex-dependent SIR effect, for which the formal housing × fetal-sex interaction analysis remained the primary statistical test.

Cross-tissue overlap was similarly limited. Nine DEGs were shared between the male fetal brain and female placenta, eight between the male fetal brain and male placenta, six between the female fetal brain and female placenta, and four between the female fetal brain and male placenta. A further seven DEGs were shared between the female fetal brain and both placental contrasts, while two were shared between the male fetal brain and both placental contrasts. No DEG was common to all four tissue–sex contrasts (Figure 2E). Thus, maternal SIR elicited a broad transcriptional response in the placenta and a more restricted response in the fetal brain. Although thresholded DEG sets differed between male and female analyses, formal interaction testing provided no evidence for significant fetal-sex differences in the SIR effect at the individual-gene level. Because limited overlap at the individual-gene level does not preclude convergence at the level of biological function, we next examined pathway and functional enrichment across the four tissue–sex contrasts.

### Pathway analysis reveals tissue- and sex-resolved responses to maternal SIR

To identify the biological processes underlying the transcriptional response to maternal SIR, we first performed Ingenuity Pathway Analysis (IPA) separately for each tissue and sex (Figure 3A-D). Pathway enrichment was quantified using Fisher’s exact test, while the IPA activation z score was used to predict pathway activation or inhibition. IPA revealed distinct functional profiles across the four tissue-sex contrasts.

**Figure 3.**
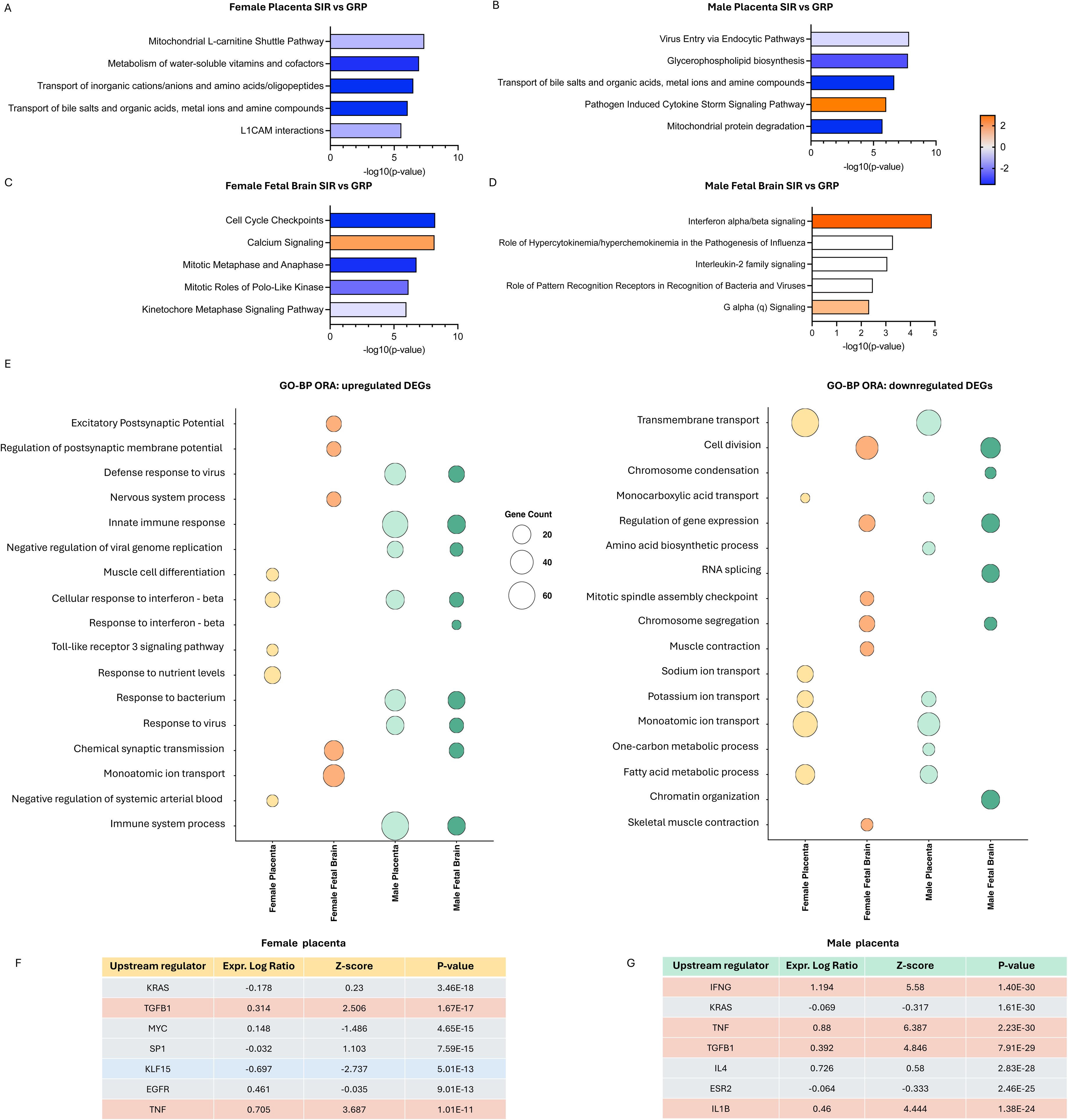
Pathway analysis reveals tissue- and sex-resolved transcriptional responses to maternal social isolation. **A–D**, Ingenuity Pathway Analysis (IPA) of DEGs identified in female placentae (**A**), male placentae (**B**), female fetal brains (**C**) and male fetal brains (**D**). The five most significantly enriched canonical pathways are shown for each tissue–sex contrast. Bar length represents pathway enrichment as −log10(*p* value), calculated using a right-tailed Fisher’s exact test. Bar colour represents the IPA activation *z score*: orange indicates predicted activation, blue indicates predicted inhibition and white indicates that no directional prediction could be assigned. **E**, Gene Ontology biological process over-representation analysis (GO-BP ORA) performed separately for upregulated and downregulated DEGs from each tissue–sex contrast. Bubble size represents the number of DEGs assigned to each biological process, and bubble colour identifies the corresponding tissue–sex contrast. Representative significantly enriched biological processes are displayed; complete enrichment results are provided in Supplementary Data 4. Enrichment was assessed relative to all expressed genes, and *p* values were corrected for multiple comparisons using the Benjamini–Hochberg procedure (*p*<0.05). **F–G**, IPA Upstream Regulator Analysis of placental DEGs from female (F) and male (G) fetuses. Selected top predicted endogenous upstream regulators are shown. Bar colour represents the IPA activation z score, with positive values indicating predicted activation and negative values indicating predicted inhibition. Complete upstream-regulator results are provided in Supplementary Data 3. GRP, group housed; SIR, social isolation rearing; DEG, differentially expressed gene; IPA, Ingenuity Pathway Analysis; GO-BP ORA, Gene Ontology biological process over-representation analysis.

In the female placenta, SIR was associated with predicted inhibition of the mitochondrial L-carnitine shuttle, water-soluble vitamin and cofactor metabolism, transport of inorganic ions and amino acids or oligopeptides, transport of bile salts and organic acids, and L1CAM interactions (Figure 3A). In the male placenta, pathogen-induced cytokine-storm signalling was predicted to be activated, whereas pathways involving viral entry, glycerophospholipid biosynthesis, solute transport and mitochondrial protein degradation were predicted to be inhibited (Figure 3B). These findings indicate that placental responses encompassed both metabolic and transport-related alterations, with additional activation of inflammatory signalling in males. Complete IPA canonical-pathway results are provided in Supplementary Data 3. Upstream-regulator analysis further supported the involvement of inflammatory signalling in the placental response to SIR (Figure 3F,G). TNF was among the predicted activated upstream regulators in both female and male placenta, with stronger predicted activation in the male placental response (activation z score = 3.69 and 6.39 in female and male placenta, respectively). The male placental response additionally showed predicted activation of inflammatory regulators including IFNG, TGFB1 and IL1B. Together with the increased placental TNFα protein concentrations, these findings provide convergent protein- and transcriptome-level support for the involvement of TNF-associated inflammatory signalling following maternal SIR. However, upstream-regulator analysis is inferential and does not establish activation of these regulators as the causal driver of the observed transcriptional changes. Complete IPA upstream-regulator results are provided in Supplementary Data 3.

Sex-stratified fetal-brain analyses highlighted different functional processes. In the female fetal brain, cell-cycle checkpoints, mitotic metaphase and anaphase, polo-like kinase functions and kinetochore signalling were predicted to be inhibited, while calcium signalling was predicted to be activated (Figure 3C). By contrast, the male fetal brain showed strong predicted activation of interferon-α/β signalling, together with enrichment of pathways involving pattern-recognition receptors, interleukin-2 family signalling, hypercytokinaemia and hyperchemokinaemia, and Gαq signalling (Figure 3D).

We next performed Gene Ontology biological process over-representation analysis (GO-BP ORA) separately for upregulated and downregulated DEGs to compare functional responses across tissues and sexes (Figure 3E). Despite the limited overlap between individual DEGs identified in the male placenta and fetal brain, their upregulated genes showed marked functional convergence on immune and antiviral processes. Both tissues were enriched for innate immune responses, defence and responses to viruses, cellular responses to interferon-β, and negative regulation of viral genome replication. The male placental response additionally included responses to bacteria, whereas the male fetal-brain response included chemical synaptic transmission. These results indicate that maternal SIR engages convergent immune and antiviral programs in the male placenta-fetal brain axis, although the underlying differentially expressed genes were largely tissue-specific.

The upregulated response in females showed less cross-tissue convergence. Female fetal-brain upregulated genes were predominantly associated with neuronal processes, including chemical synaptic transmission, regulation of postsynaptic membrane potential and excitatory postsynaptic potential. Upregulated genes in the female placenta instead included processes related to nutrient sensing, Toll-like receptor 3 signalling and muscle-cell differentiation.

A different pattern emerged among downregulated genes. Here, the strongest functional similarity was observed between male and female placentae, both of which showed enrichment for transmembrane transport, monoatomic and potassium-ion transport, monocarboxylic acid transport, and fatty-acid metabolism. Female placental downregulated genes were additionally enriched for sodium-ion transport, whereas the male placental response included amino-acid biosynthesis and one-carbon metabolism. Thus, although the individual placental DEG profiles were not identical, downregulated genes converged on alterations in placental transport and metabolic functions across sexes.

In contrast, the downregulated fetal-brain profiles were more divergent. Female fetal-brain downregulated genes were strongly enriched for cell division, chromosome segregation and mitotic-spindle checkpoint processes. Downregulated genes in the male fetal brain were primarily associated with regulation of gene expression, RNA splicing, chromatin organization and chromosome condensation, although cell-division and chromosome-segregation processes were also represented. Together, these analyses reveal direction-dependent patterns of functional convergence: upregulated genes link the male placenta and fetal brain through immune and antiviral signalling, whereas downregulated genes reveal shared transport and metabolic alterations across male and female placentae. Complete GO-BP enrichment results are provided in Supplementary Data 4.

### SIR-responsive transcriptional networks reveal cross-tissue associations that are largely explained by maternal housing

To determine whether transcriptional programmes responsive to maternal SIR covaried across the placenta and fetal brain, we first performed weighted gene co-expression network analysis (WGCNA) using genes differentially expressed following SIR. DEGs were organized into co-expression modules independently within placenta and fetal brain, and module eigengenes were correlated across matched tissues. Analyses were performed using all DEGs and separately for upregulated and downregulated genes, across the complete dataset and within male and female fetuses.

The initial analysis identified extensive cross-tissue covariation among SIR-responsive modules, with 22 of 27 tested placenta–brain module pairs showing significant correlations after multiple-testing correction (Figure 4A). Because these networks were constructed from genes selected according to their response to SIR, however, the observed correlations could reflect the shared effect of maternal housing on both tissues. We therefore reassessed these associations after adjustment for housing and using a litter-aware permutation procedure. Housing adjustment markedly attenuated the DEG-derived correlations, reducing the mean absolute correlation coefficient from 0.544 to 0.258, with only one of the 22 initially significant associations remaining at FDR < 0.10 (Figure 4B). This indicated that much of the apparent cross-tissue covariation among SIR-responsive modules was attributable to their shared association with maternal housing.

**Figure 4.**
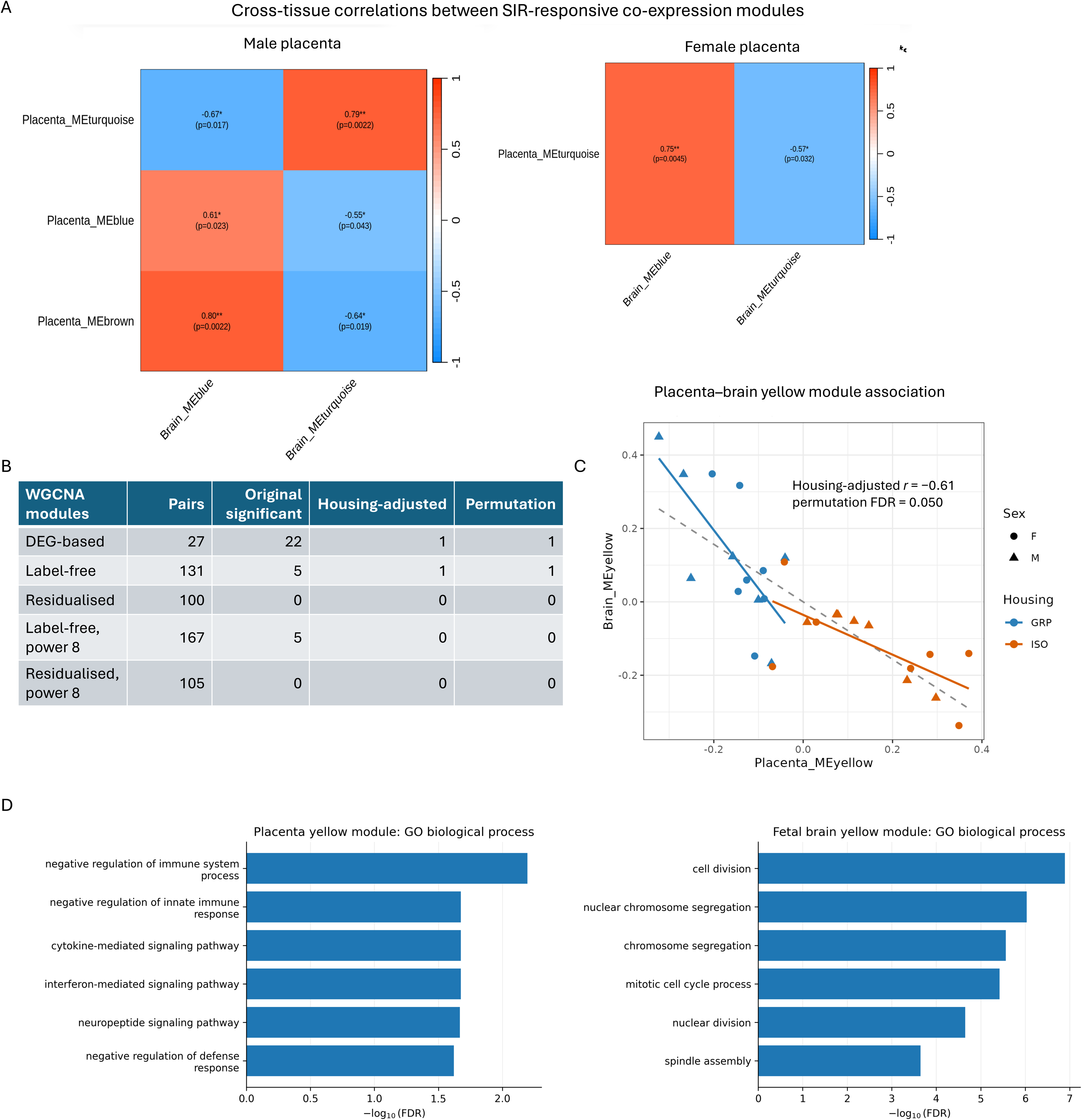
Cross-tissue co-expression analysis identifies an exploratory association between placental immune and fetal-brain proliferative programmes. **A**, Cross-tissue correlations between module eigengenes derived from SIR-responsive differentially expressed genes in male and female placenta–fetal brain pairs. Values indicate Pearson correlation coefficients, with corresponding adjusted *P* values shown in parentheses. **B**, Summary of cross-tissue module associations across WGCNA approaches. DEG-based networks were constructed from SIR-responsive genes; exposure-independent networks were constructed from the 5,000 most variably expressed genes independently of housing; residualised networks were constructed after removal of housing-associated variation. Sensitivity analyses using the recommended soft-thresholding power of 8 are also shown. Numbers indicate the total placenta–brain module pairs tested and the number meeting the indicated multiple-testing thresholds before and after adjustment for maternal housing. **C**, Association between placental and fetal-brain yellow-module eigengenes identified in the exposure-independent WGCNA across all 28 matched samples. Points represent individual fetuses, with colour indicating maternal housing condition and shape indicating fetal sex. Solid lines indicate associations within GRP and SIR groups and the dashed line indicates the pooled association. The housing-adjusted correlation and litter-aware permutation FDR are shown. **D**, Gene Ontology Biological Process enrichment of genes belonging to the placental and fetal-brain yellow modules. Bars show −log10(FDR) for representative significantly enriched biological processes. GRP, group housed; SIR, social isolation rearing; WGCNA, weighted gene co-expression network analysis; FDR, false discovery rate. Complete WGCNA module assignments and cross-tissue correlation results are provided in Supplementary Data 5.

We next asked whether placenta–brain associations could be detected without selecting genes according to their response to SIR. Independent co-expression networks were therefore constructed from the 5,000 most variably expressed genes in each tissue, irrespective of maternal housing. Of 131 placenta–brain module pairs, five were significant before adjustment and one remained following adjustment for maternal housing and litter-aware permutation testing (Figure 4B,C). This association linked the placental yellow module with the fetal-brain yellow module and was inverse in direction (pooled r = −0.78; housing-adjusted r = −0.61; conventional FDR = 0.040; permutation FDR = 0.050). The association was similar within GRP (r = −0.70) and SIR (r = −0.70) pregnancies and remained comparable after additional adjustment for fetal sex (r = −0.60).

Functional enrichment revealed distinct biological programmes within the associated modules (Figure 4D). The placental yellow module was enriched for immune-related processes, including interferon- and cytokine-mediated signalling and regulation of innate immune responses, whereas the fetal-brain yellow module showed strong enrichment for cell division, chromosome segregation and mitotic cell-cycle processes. The placental module was positively associated with SIR, whereas the fetal-brain module was negatively associated with SIR. Further analyses indicated that the inverse placenta–brain relationship was predominantly evident at the pregnancy level: litter-averaged placental and fetal-brain module eigengenes were negatively correlated across pregnancies (r = −0.67, p = 0.012), whereas within-litter male–female differences were not correlated (r = −0.18, p = 0.53). Together, these analyses identify an exploratory cross-tissue association between a placental immune-related transcriptional programme and a fetal-brain proliferative programme, while indicating that the broader correlations observed among SIR-responsive networks are largely explained by their common relationship with maternal housing. Complete module assignments, cross-tissue module-correlation results and functional-enrichment results are provided in Supplementary Data 5.

## Discussion

In this study, we investigated how prolonged social adversity initiated before conception and maintained throughout pregnancy alters molecular signalling across the placenta–fetal brain axis. Maternal SIR produced inflammatory alterations in both placenta and fetal brain and a broad transcriptional response that was substantially more extensive in the placenta than in the developing brain. Although individual differentially expressed genes showed relatively little overlap across tissues, functional analyses identified convergence on immune and antiviral signalling, placental transport and metabolic processes, and fetal-brain proliferative and neuronal programmes. Importantly, formal interaction analyses provided no evidence that the transcriptional response to SIR differed significantly by fetal sex at the individual-gene level, despite differences between male- and female-stratified DEG and pathway profiles. Finally, cross-tissue network analyses showed that much of the apparent covariation between SIR-responsive placental and fetal-brain modules could be attributed to their shared response to maternal housing. An unbiased analysis nevertheless identified an inverse pregnancy-level association between a placental immune-related module and a fetal-brain module enriched for cell-cycle and proliferative processes. Together, these findings indicate that prolonged maternal social adversity is biologically embedded across both sides of the placenta–fetal brain interface, while highlighting the distinction between parallel responses to a common maternal exposure and direct molecular coupling between the two tissues.

### Inflammatory alterations across placenta and fetal brain

A prominent finding was the presence of inflammatory alterations in both placenta and fetal brain at late gestation. SIR increased placental TNFα and IL-6, with a trend towards increased IFNγ, while fetal brains showed increased IL-6 and IL-1β but reduced IFNγ. This pattern is consistent with evidence that psychological stress during pregnancy can generate an inflammatory intrauterine environment without an overt maternal immune challenge. In humans, antenatal and perinatal depression has been associated with altered inflammatory physiology, although findings vary considerably across cohorts and gestational time points. Depressive symptoms during pregnancy have been associated with elevated circulating IL-6 and, in some studies, TNFα, while clinically diagnosed antenatal depression has similarly been associated with increased IL-6, TNFα and other inflammatory mediators^9,10,15^. More recently, a systematic review of 56 studies found that the most frequently reported associations involved increased IL-6, TNFα, CRP and IL-8, while also highlighting substantial heterogeneity in biomarker sampling, timing and depression assessment^46^. This pattern is consistent with the broader depression literature, in which elevations of IL-6, CRP and TNFα are among the most reproducible inflammatory alterations^47,48^.

Experimental studies provide stronger evidence that psychological stress itself can alter inflammatory signalling within the intrauterine compartment. Late-gestational restraint stress has been shown to induce inflammation in both placenta and fetal brain, including increased CCL2 in both tissues and CCL2-dependent elevation of fetal-brain IL-6^49^. These effects were strongly influenced by the maternal microbiota, indicating that stress-induced intrauterine inflammation can emerge through interactions between maternal physiological systems^49^. Other restraint-stress studies have similarly identified alterations in cytokines across maternal, placental and fetal compartments, although the specific mediators and direction of change vary according to tissue and gestational timing^50^. Comparative analyses of different prenatal-stress paradigms further indicate that placental inflammatory responses depend strongly on the nature and timing of the maternal exposure, with distinct effects on placental IL-6, TNFα, IL-1β and related immune pathways across stress models^51^. Together, these studies support an intrauterine inflammatory response to psychological stress that varies according to tissue, timing and stress paradigm, consistent with the distinct placental and fetal-brain cytokine profiles observed here.

The increase in IL-6 in both placenta and fetal brain is particularly notable given experimental evidence that IL-6 signalling at the maternal–placental interface can influence fetal-brain inflammatory responses and neurodevelopment. In a maternal immune-activation model, trophoblast-specific deletion of the IL-6 receptor prevented MIA-induced inflammatory responses in both placenta and fetal brain and attenuated subsequent neuropathological and behavioural abnormalities in the offspring^52^. Although maternal immune activation represents a mechanistically distinct and more acute inflammatory exposure than the chronic social adversity modelled here, this provides causal evidence that placental IL-6 signalling can participate in communication between maternal inflammation and the developing brain. Our data do not establish such a causal pathway, but the parallel increase in IL-6 across placenta and fetal brain identifies IL-6 signalling as one potential component of the response to SIR.

At the same time, the cytokine profiles were not simply mirrored across tissues: TNFα was increased in placenta but not fetal brain, whereas IL-1β was increased in fetal brain but not placenta, and IFNγ showed opposite directional patterns. This argues against interpreting the fetal inflammatory phenotype as passive transmission of placental cytokine concentrations. Rather, maternal adversity appears to engage tissue-specific immune responses across the maternal–placental–fetal system. The placenta may therefore act as a responsive immune interface, potentially modifying maternal inflammatory signals or generating secondary signals that influence fetal tissues. However, causal manipulation will be required to determine whether placental inflammatory signalling contributes directly to the fetal-brain response.

### Maternal SIR produces broad placental and more restricted fetal-brain transcriptional responses

The transcriptomic findings reinforce the placenta as a major site of adaptation to maternal adversity. SIR altered more than 1,000 placental transcripts in each sex-stratified analysis, compared with approximately 100–130 transcripts in fetal brain. Such a difference is consistent with the placenta’s position at the maternal–fetal interface, where it responds dynamically to maternal metabolic, endocrine and immune conditions while supporting fetal homeostasis and development^53,54^. Human placental gene-expression studies similarly indicate that maternal psychological adversity is associated with molecular alterations in the placenta. In a large RNA-sequencing study of 1,029 mother–child pairs, Baker et al.^41^ identified placental transcriptional signatures associated with both prenatal stressful life events and preconceptional maternal adversity, including alterations in endoplasmic-reticulum protein processing, protein synthesis and metabolic pathways. Earlier studies have likewise reported altered placental expression of genes involved in nutrient sensing and transport, hypoxia responses, glucocorticoid signalling and fetal growth in association with maternal stress, depression or anxiety ^55,56^. Studies specifically examining antenatal depression have also identified broader placental gene-expression differences, although findings have varied between cohorts and separating the effects of maternal depression from those of antidepressant exposure remains challenging^57^.

The functional profile of the placental response further reflected its combined immune and metabolic/transport functions. Male-stratified placental DEGs showed prominent immune and antiviral signatures, including predicted cytokine-storm signalling and enrichment of interferon-related processes, whereas downregulated placental genes in both sex-stratified analyses converged on transmembrane and ion transport, fatty-acid metabolism and related metabolic processes. These findings broadly parallel human studies linking maternal psychological adversity with placental pathways involved in cellular stress and metabolism, as well as altered expression of genes involved in nutrient sensing and transport, including OGT and GLUT1^55^. The convergence between increased placental TNFα protein and predicted activation of TNF as an upstream regulator in both sex-stratified placental transcriptomic analyses is particularly notable, although upstream-regulator analysis is inferential and does not establish TNF as the causal driver of the transcriptional response.

The fetal-brain transcriptional response was quantitatively smaller but implicated processes directly relevant to neurodevelopment. Female-stratified analyses highlighted reduced cell-cycle, chromosome-segregation and mitotic programmes, whereas male-stratified analyses showed prominent interferon and innate immune signalling. Prenatal-stress transcriptomic studies have independently implicated both classes of processes. Whole fetal-brain RNA sequencing following chronic unpredictable maternal stress has identified alterations in cell-cycle and neurodevelopmental programmes, with cell-cycle-associated transcripts particularly enriched in proliferative radial-glial populations and neuronal/synaptic genes showing distinct responses^58^. Although the direction of the cell-cycle effect in that model was not identical to that observed here, the convergence on proliferative programmes suggests that regulation of neural progenitor dynamics may be a recurring target of maternal stress, with direction and magnitude likely depending on developmental stage and stress paradigm. A recent study analysing matched E18.5 placenta and fetal brain after prenatal stress identified interferon-β-related transcriptional alterations in both tissues, although in that paradigm IFN-β signalling was suppressed rather than activated^59^. These differences emphasize that “prenatal stress” does not constitute a single biological exposure; timing, chronicity, stressor type and maternal physiological response are likely to determine the molecular state ultimately encountered by the fetus.

Notably, convergence across placenta and fetal brain was more evident at the level of biological function than at the level of individual genes. Relatively few DEGs were shared directly between tissues, yet male-stratified upregulated genes in both placenta and fetal brain converged strongly on innate immune, antiviral and interferon-related processes. This suggests that a common maternal perturbation need not produce identical transcriptional responses in tissues with profoundly different cellular composition and biological functions. Instead, different gene sets may engage related biological programmes. This distinction is important when considering the placenta as a potential molecular readout of the intrauterine environment: placental transcriptional changes should not necessarily be expected to reproduce fetal-brain gene-expression changes gene-for-gene.

### Sex-stratified responses do not provide evidence for statistically significant sex-dependent effects

Fetal sex is widely proposed to modify placental responses to maternal adversity, and previous studies have reported divergent molecular responses in male and female placenta and fetal brain following prenatal and preconceptional stress^38,60–62^. Cissé et al., for example, reported markedly different pathway-level responses in male and female placenta and fetal brain following parental preconception stress, while subsequent work from the same group identified divergent placental metabolic, transport and immune programmes between male and female offspring^38,60^. Human studies have likewise reported different associations between maternal stress and placental gene expression when analyses are stratified by fetal sex^55^. Similarly, in our sex-stratified analyses, immune and antiviral signalling was particularly prominent in male placenta and fetal brain, whereas female fetal-brain DEGs highlighted cell-cycle and neuronal processes. However, formal maternal housing × fetal-sex interaction testing identified no individual transcript for which the SIR effect differed significantly between sexes after correction for multiple testing. Similarly, cytokine concentrations showed no significant housing × sex interactions. This distinction is important because significance in one sex but not the other does not itself demonstrate a significant difference between sexes. Consequently, our results support fetal sex as an informative stratification variable but do not provide statistical evidence for gene-level sex-specific responses to SIR.

This finding does not exclude more subtle sex modulation. Interaction tests are generally less powerful than tests of a main exposure effect, and the present study included seven pregnancies per housing condition. Moreover, sex-dependent responses may emerge at the level of cell composition, pathway organization, developmental trajectory or later phenotype without producing large individual-gene interaction effects at GD18. Recent experimental work illustrates the value of explicitly testing these interactions: comparisons of multiple prenatal-stress paradigms have identified stress × sex interactions for selected placental immune genes while finding many other effects shared across sexes^51^. Thus, rather than classifying molecular effects as sex-specific on the basis of separate male and female significance tests, future adequately powered studies should distinguish shared exposure effects from formally supported sex-dependent responses.

### Cross-tissue network analyses distinguish common exposure effects from placenta–brain covariation

A central aim of this study was to determine whether molecular responses in placenta and fetal brain covaried within pregnancy. Initial WGCNA of SIR-responsive genes suggested extensive cross-tissue coordination, with 22 of 27 module pairs significantly correlated. However, these networks were constructed from genes selected precisely because they responded to SIR. Once the shared effect of maternal housing was accounted for, almost all of these associations disappeared. This highlights an important distinction: two tissues can show highly coordinated responses at the group level because both respond to the same maternal environment without variation in one tissue necessarily tracking variation in the other within pregnancies.

Previous work supports the possibility that placental and fetal-brain transcriptional states are related. Network analyses in normally developing mouse pregnancies have identified correlated transcriptional architecture between placenta and fetal brain, providing evidence for coordinated molecular programmes across the two tissues^63^. Our analyses extend this work by examining cross-tissue transcriptional covariance following maternal adversity and by explicitly separating correlations attributable to the experimental exposure from residual cross-pregnancy covariation. When networks were constructed independently of the SIR response, one cross-tissue association remained after adjustment for housing and litter-aware permutation testing. Higher expression of a placental module enriched for immune, cytokine and interferon-related processes was associated with lower expression of a fetal-brain module enriched for cell division, chromosome segregation and mitotic processes. Both components of this relationship therefore recapitulated biological themes independently identified in the differential-expression analyses. The association is intriguing in light of experimental evidence that inflammatory cytokines can directly alter neural progenitor proliferation and differentiation^64^ and that prenatal stress can reduce neural progenitor proliferation during fetal brain development^65,66^. However, the association should remain explicitly exploratory. It does not demonstrate that placental immune activation suppresses fetal-brain proliferation, nor does it establish the placenta as the mediator of the relationship. Maternal endocrine, metabolic, immune or microbial factors could independently influence both tissues.

Indeed, the association was predominantly expressed at the pregnancy level, rather than through male–female variation within individual litters. This is biologically informative: it suggests that pregnancies characterized by a relatively stronger placental immune programme also tended to show a relatively weaker fetal-brain proliferative programme. Such a pattern is compatible with a shared maternal physiological state, placenta-mediated signalling, or both. Determining among these possibilities will require causal experiments in which candidate placental pathways are manipulated independently of the maternal exposure.

### Strengths, limitations and implications

Several aspects of the study strengthen interpretation of these findings. Placenta and fetal brain were obtained as matched samples from the same fetuses, while the inclusion of one male and one female fetus from each pregnancy enabled litter structure to be incorporated explicitly into both interaction and cross-tissue analyses. The combination of cytokine measurements with transcriptome-wide profiling further allowed inflammatory alterations to be considered alongside broader molecular programmes. Finally, the use of both exposure-selected and exposure-independent WGCNA illustrates the importance of distinguishing common experimental effects from genuine cross-sample covariation.

The study also has limitations. The number of independent pregnancies was modest, particularly for detecting housing × fetal-sex interactions, and bulk RNA sequencing cannot distinguish transcriptional regulation within individual cell types from shifts in placental or fetal-brain cellular composition. The whole fetal brain additionally contains multiple regions and developmental cell populations that may respond differently to maternal adversity. Maternal SIR begins before conception and continues throughout gestation, meaning that the present design cannot determine whether the observed fetal changes originate from preconceptional programming, the maternal state during pregnancy, or their combination. Finally, the cross-tissue analyses are correlational and cannot establish the direction or mechanism of placenta–brain communication.

Overall, our findings show that prolonged maternal social adversity beginning before conception alters inflammatory and transcriptional states across both placenta and developing fetal brain. The placenta exhibited a particularly broad response encompassing inflammatory, metabolic and transport-related programmes, whereas fetal-brain alterations implicated immune, neuronal and proliferative processes. Despite descriptively different male and female profiles, formal interaction analyses did not support robust gene-level sex-specific responses. Finally, cross-tissue network analysis suggests that much of the apparent coordination between placenta and fetal brain reflects their common response to the maternal environment, while identifying an exploratory pregnancy-level relationship between placental immune signalling and fetal-brain proliferative programmes. These findings position the placenta not simply as a surrogate for fetal-brain molecular state, but as a responsive component of a broader maternal–placental–fetal system whose contribution to fetal neurodevelopment now requires causal investigation.

## Data availability

The datasets generated and analysed during the current study are available from the corresponding author upon reasonable request. The RNA-sequencing data are not publicly deposited at the time of preprint publication and will be made available in accordance with journal data-sharing requirements upon publication of the final peer-reviewed article.

## Author contributions

J.R. conceived and supervised the study and acquired funding. M.I. performed the experiments and contributed to data collection and analysis. J.R.R. performed bioinformatic and statistical analyses. F.H. assisted with the in vivo experiments. R.S. assisted with the in vivo experiments. A.C. provided mentorship. M.I., J.R.and J.R.R. interpreted the data and drafted the manuscript. All authors contributed to critical revision of the manuscript and approved the final version.

## References

1 Fisher, J., Cabral de Mello, M., Patel, V., Rahman, A., Tran, T., Holton, S., et al. Prevalence and determinants of common perinatal mental disorders in women in low- and lower-middle-income countries: a systematic review. Bull World Health Organ 90, 139G–149G, doi:10.2471/BLT.11.091850 (2012).

2 Howard, L. M., Molyneaux, E., Dennis, C. L., Rochat, T., Stein, A. & Milgrom, J. Non-psychotic mental disorders in the perinatal period. Lancet 384, 1775–1788, doi:10.1016/S0140-6736(14)61276-9 (2014).

3 Melville, J. L., Gavin, A., Guo, Y., Fan, M. Y. & Katon, W. J. Depressive disorders during pregnancy: prevalence and risk factors in a large urban sample. Obstet Gynecol 116, 1064–1070, doi:10.1097/AOG.0b013e3181f60b0a (2010).

4 Yin, X., Sun, N., Jiang, N., Xu, X., Gan, Y., Zhang, J. et al. Prevalence and associated factors of antenatal depression: Systematic reviews and meta-analyses. Clin Psychol Rev 83, 101932, doi:10.1016/j.cpr.2020.101932 (2021).

5 Hay, D. F., Pawlby, S., Waters, C. S. & Sharp, D. Antepartum and postpartum exposure to maternal depression: different effects on different adolescent outcomes. J Child Psychol Psychiatry 49, 1079–1088, doi:10.1111/j.1469-7610.2008.01959.x (2008).

6 Lahti, M., Savolainen, K., Tuovinen, S., Pesonen, A. K., Lahti, J., Heinonen, K. et al. Maternal Depressive Symptoms During and After Pregnancy and Psychiatric Problems in Children. J Am Acad Child Adolesc Psychiatry 56, 30–39 e37, doi:10.1016/j.jaac.2016.10.007 (2017).

7 Tuovinen, S., Lahti-Pulkkinen, M., Girchenko, P., Heinonen, K., Lahti, J., Reynolds, R. M. et al. Maternal antenatal stress and mental and behavioral disorders in their children. J Affect Disord 278, 57–65, doi:10.1016/j.jad.2020.09.063 (2021).

8 Leff Gelman, P., Mancilla-Herrera, I., Flores-Ramos, M., Saravia Takashima, M. F., Cruz Coronel, F. M., Cruz Fuentes, C., et al. The cytokine profile of women with severe anxiety and depression during pregnancy. BMC Psychiatry 19, 104, doi:10.1186/s12888-019-2087-6 (2019).

9 Leff-Gelman, P., Mancilla-Herrera, I., Flores-Ramos, M., Cruz-Fuentes, C., Reyes-Grajeda, J. P., Garcia-Cuetara Mdel, P. et al. The Immune System and the Role of Inflammation in Perinatal Depression. Neurosci Bull 32, 398–420, doi:10.1007/s12264-016-0048-3 (2016).

10 Osborne, S., Biaggi, A., Chua, T. E., Du Preez, A., Hazelgrove, K., Nikkheslat, N. et al. Antenatal depression programs cortisol stress reactivity in offspring through increased maternal inflammation and cortisol in pregnancy: The Psychiatry Research and Motherhood - Depression (PRAM-D) Study. Psychoneuroendocrinology 98, 211–221, doi:10.1016/j.psyneuen.2018.06.017 (2018).

11 Sha, Q., Madaj, Z., Keaton, S., Escobar Galvis, M. L., Smart, L., Krzyzanowski, S. et al. Cytokines and tryptophan metabolites can predict depressive symptoms in pregnancy. Transl Psychiatry 12, 35, doi:10.1038/s41398-022-01801-8 (2022).

12 Maes, M., Abe, Y., Sirichokchatchawan, W., Suwimonteerabutr, J., Sangkomkamhangd, U., Almulla, A. F. et al. The Cytokine, Chemokine, and Growth Factor Network of Prenatal Depression. Brain Sci 13, doi:10.3390/brainsci13050727 (2023).

13 Serati, M., Redaelli, M., Buoli, M. & Altamura, A. C. Perinatal Major Depression Biomarkers: A systematic review. J Affect Disord 193, 391–404, doi:10.1016/j.jad.2016.01.027 (2016).

14 Cassidy-Bushrow, A. E., Burmeister, C., Birbeck, J., Chen, Y., Lamerato, L., Lemke, L. D. et al. Ambient BTEX exposure and mid-pregnancy inflammatory biomarkers in pregnant African American women. J Reprod Immunol 145, 103305, doi:10.1016/j.jri.2021.103305 (2021).

15 Christian, L. M., Franco, A., Glaser, R. & Iams, J. D. Depressive symptoms are associated with elevated serum proinflammatory cytokines among pregnant women. Brain Behav Immun 23, 750–754, doi:10.1016/j.bbi.2009.02.012 (2009).

16 Lahti-Pulkkinen, M., Girchenko, P., Robinson, R., Lehto, S. M., Toffol, E., Heinonen, K. et al. Maternal depression and inflammation during pregnancy. Psychol Med 50, 1839–1851, doi:10.1017/S0033291719001909 (2020).

17 Naude, P. J. W., Pariante, C., Hoffman, N., Koopowitz, S. M., Donald, K. A., Zar, H. J. et al. Antenatal maternal depression, early life inflammation and neurodevelopment in a South African birth cohort. Brain Behav Immun 105, 160–168, doi:10.1016/j.bbi.2022.07.001 (2022).

18 Janssen, A. B., Kertes, D. A., McNamara, G. I., Braithwaite, E. C., Creeth, H. D., Glover, V. I. et al. A Role for the Placenta in Programming Maternal Mood and Childhood Behavioural Disorders. J Neuroendocrinol 28, n/a, doi:10.1111/jne.12373 (2016).

19 Bronson, S. L. & Bale, T. L. Prenatal stress-induced increases in placental inflammation and offspring hyperactivity are male-specific and ameliorated by maternal antiinflammatory treatment. Endocrinology 155, 2635–2646, doi:10.1210/en.2014-1040 (2014).

20 Rakers, F., Rupprecht, S., Dreiling, M., Bergmeier, C., Witte, O. W. & Schwab, M. Transfer of maternal psychosocial stress to the fetus. Neurosci Biobehav Rev, doi:10.1016/j.neubiorev.2017.02.019 (2017).

21 Scott, H., Phillips, T. J., Sze, Y., Alfieri, A., Rogers, M. F., Volpato, V. et al. Maternal antioxidant treatment prevents the adverse effects of prenatal stress on the offspring’s brain and behavior. Neurobiol Stress 13, 100281, doi:10.1016/j.ynstr.2020.100281 (2020).

22 Ding, J., Maxwell, A., Adzibolosu, N., Hu, A., You, Y., Liao, A. et al. Mechanisms of immune regulation by the placenta: Role of type I interferon and interferon-stimulated genes signaling during pregnancy. Immunol Rev 308, 9–24, doi:10.1111/imr.13077 (2022).

23 Han, V. X., Patel, S., Jones, H. F., Nielsen, T. C., Mohammad, S. S., Hofer, M. J. et al. Maternal acute and chronic inflammation in pregnancy is associated with common neurodevelopmental disorders: a systematic review. Transl Psychiatry 11, 71, doi:10.1038/s41398-021-01198-w (2021).

24 Kirkpatrick, M., Mandal, G., Elhadidy, I., Mariani, N., Priestley, K., Pariante, C. M. et al. From placenta to the foetus: a systematic review of in vitro models of stress-and inflammation-induced depression in pregnancy. Mol Psychiatry, doi:10.1038/s41380-024-02866-1 (2024).

25 Graham, A. M., Doyle, O., Tilden, E. L., Sullivan, E. L., Gustafsson, H. C., Marr, M. et al. Effects of Maternal Psychological Stress During Pregnancy on Offspring Brain Development: Considering the Role of Inflammation and Potential for Preventive Intervention. Biol Psychiatry Cogn Neurosci Neuroimaging 7, 461–470, doi:10.1016/j.bpsc.2021.10.012 (2022).

26 Raikkonen, K., Pesonen, A. K., O’Reilly, J. R., Tuovinen, S., Lahti, M., Kajantie, E. et al. Maternal depressive symptoms during pregnancy, placental expression of genes regulating glucocorticoid and serotonin function and infant regulatory behaviors. Psychol Med 45, 3217–3226, doi:10.1017/S003329171500121X (2015).

27 Capron, L. E., Ramchandani, P. G. & Glover, V. Maternal prenatal stress and placental gene expression of NR3C1 and HSD11B2: The effects of maternal ethnicity. Psychoneuroendocrinology 87, 166–172, doi:10.1016/j.psyneuen.2017.10.019 (2018).

28 Edvinsson, A., Hellgren, C., Kunovac Kallak, T., Akerud, H., Skalkidou, A., Stener-Victorin, E. et al. The effect of antenatal depression and antidepressant treatment on placental tissue: a protein-validated gene expression study. BMC Pregnancy Childbirth 19, 479, doi:10.1186/s12884-019-2586-y (2019).

29 Martinez, C. A., Marteinsdottir, I., Josefsson, A., Sydsjo, G., Theodorsson, E. & Rodriguez-Martinez, H. Prenatal stress, anxiety and depression alter transcripts, proteins and pathways associated with immune responses at the maternal-fetal interfacedagger. Biol Reprod 106, 449–462, doi:10.1093/biolre/ioab232 (2022).

30 Ponder, K. L., Salisbury, A., McGonnigal, B., Laliberte, A., Lester, B. & Padbury, J. F. Maternal depression and anxiety are associated with altered gene expression in the human placenta without modification by antidepressant use: implications for fetal programming. Dev Psychobiol 53, 711–723, doi:10.1002/dev.20549 (2011).

31 Reynolds, R. M., Pesonen, A. K., O’Reilly, J. R., Tuovinen, S., Lahti, M., Kajantie, E. et al. Maternal depressive symptoms throughout pregnancy are associated with increased placental glucocorticoid sensitivity. Psychol Med 45, 2023–2030, doi:10.1017/S003329171400316X (2015).

32 Boksa, P. Effects of prenatal infection on brain development and behavior: a review of findings from animal models. Brain Behav Immun 24, 881–897, doi:10.1016/j.bbi.2010.03.005 (2010).

33 Maccari, S. & Morley-Fletcher, S. Effects of prenatal restraint stress on the hypothalamus-pituitary-adrenal axis and related behavioural and neurobiological alterations. Psychoneuroendocrinology 32 **Suppl 1**, S10–15, doi:10.1016/j.psyneuen.2007.06.005 (2007).

34 Creutzberg, K. C., Sanson, A., Viola, T. W., Marchisella, F., Begni, V., Grassi-Oliveira, R. et al. Long-lasting effects of prenatal stress on HPA axis and inflammation: A systematic review and multilevel meta-analysis in rodent studies. Neurosci Biobehav Rev 127, 270–283, doi:10.1016/j.neubiorev.2021.04.032 (2021).

35 Enayati, M., Mosaferi, B., Homberg, J. R., Diniz, D. M. & Salari, A. A. Prenatal maternal stress alters depression-related symptoms in a strain - and sex-dependent manner in rodent offspring. Life Sci 251, 117597, doi:10.1016/j.lfs.2020.117597 (2020).

36 Weinstock, M. Prenatal stressors in rodents: Effects on behavior. Neurobiol Stress 6, 3–13, doi:10.1016/j.ynstr.2016.08.004 (2017).

37 Biaggi, A., Conroy, S., Pawlby, S. & Pariante, C. M. Identifying the women at risk of antenatal anxiety and depression: A systematic review. J Affect Disord 191, 62–77, doi:10.1016/j.jad.2015.11.014 (2016).

38 Cisse, Y. M., Chan, J. C., Nugent, B. M., Banducci, C. & Bale, T. L. Brain and placental transcriptional responses as a readout of maternal and paternal preconception stress are fetal sex specific. Placenta 100, 164–170, doi:10.1016/j.placenta.2020.06.019 (2020).

39 Gonzalez, T. L., Sun, T., Koeppel, A. F., Lee, B., Wang, E. T., Farber, C. R. et al. Sex differences in the late first trimester human placenta transcriptome. Biol Sex Differ 9, 4, doi:10.1186/s13293-018-0165-y (2018).

40 Braun, A. E., Mitchel, O. R., Gonzalez, T. L., Sun, T., Flowers, A. E., Pisarska, M. D. et al. Sex at the interface: the origin and impact of sex differences in the developing human placenta. Biol Sex Differ 13, 50, doi:10.1186/s13293-022-00459-7 (2022).

41 Baker, B. H., Freije, S., MacDonald, J. W., Bammler, T. K., Benson, C., Carroll, K. N. et al. Placental transcriptomic signatures of prenatal and preconceptional maternal stress. Mol Psychiatry 29, 1179–1191, doi:10.1038/s41380-023-02403-6 (2024).

42 Scarborough, J., Mueller, F. S., Weber-Stadlbauer, U., Mattei, D., Opitz, L., Cattaneo, A. et al. A novel murine model to study the impact of maternal depression and antidepressant treatment on biobehavioral functions in the offspring. Mol Psychiatry 26, 6756–6772, doi:10.1038/s41380-021-01145-7 (2021).

43 Scarborough, J., Iachizzi, M., Schalbetter, S. M., Muller, F. S., Weber-Stadlbauer, U. & Richetto, J. Prenatal and postnatal influences on behavioral development in a mouse model of preconceptional stress. Neurobiol Stress 29, 100614, doi:10.1016/j.ynstr.2024.100614 (2024).

44 Mueller, F. S., Richetto, J., Hayes, L. N., Zambon, A., Pollak, D. D., Sawa, A. et al. Influence of poly(I:C) variability on thermoregulation, immune responses and pregnancy outcomes in mouse models of maternal immune activation. Brain Behav Immun 80, 406–418, doi:10.1016/j.bbi.2019.04.019 (2019).

45 Mueller, F. S., Polesel, M., Richetto, J., Meyer, U. & Weber-Stadlbauer, U. Mouse models of maternal immune activation: Mind your caging system! Brain Behav Immun 73, 643–660, doi:10.1016/j.bbi.2018.07.014 (2018).

46 Silva-Fernandes, A., Conde, A., Marques, M., Caparros-Gonzalez, R. A., Fransson, E., Mesquita, A. R. et al. Inflammatory biomarkers and perinatal depression: A systematic review. PLoS One 19, e0280612, doi:10.1371/journal.pone.0280612 (2024).

47 Haapakoski, R., Mathieu, J., Ebmeier, K. P., Alenius, H. & Kivimaki, M. Cumulative meta-analysis of interleukins 6 and 1beta, tumour necrosis factor alpha and C-reactive protein in patients with major depressive disorder. Brain Behav Immun 49, 206–215, doi:10.1016/j.bbi.2015.06.001 (2015).

48 Osimo, E. F., Pillinger, T., Rodriguez, I. M., Khandaker, G. M., Pariante, C. M. & Howes, O. D. Inflammatory markers in depression: A meta-analysis of mean differences and variability in 5,166 patients and 5,083 controls. Brain Behav Immun 87, 901–909, doi:10.1016/j.bbi.2020.02.010 (2020).

49 Chen, H. J., Antonson, A. M., Rajasekera, T. A., Patterson, J. M., Bailey, M. T. & Gur, T. L. Prenatal stress causes intrauterine inflammation and serotonergic dysfunction, and long-term behavioral deficits through microbe- and CCL2-dependent mechanisms. Transl Psychiatry 10, 191, doi:10.1038/s41398-020-00876-5 (2020).

50 Gumusoglu, S. B., Maurer, S. V. & Stevens, H. E. Dataset describing maternal prenatal restraint stress effects on immune factors in mice. Data Brief 43, 108348, doi:10.1016/j.dib.2022.108348 (2022).

51 Dukle, M., Fairbairn, F. M., Maurer, S. V., Grueter, M. M., Baig, A. I., Telles, R. et al. Comparative Analysis of Prenatal Stress Models: Placental and Neurodevelopmental Outcomes in Mice. Yale J Biol Med 99, 127–141, doi:10.59249/MHFP3728 (2026).

52 Wu, W. L., Hsiao, E. Y., Yan, Z., Mazmanian, S. K. & Patterson, P. H. The placental interleukin-6 signaling controls fetal brain development and behavior. Brain Behav Immun 62, 11–23, doi:10.1016/j.bbi.2016.11.007 (2017).

53 Semmes, E. C. & Coyne, C. B. Innate immune defenses at the maternal-fetal interface. Curr Opin Immunol 74, 60–67, doi:10.1016/j.coi.2021.10.007 (2022).

54 O’Brien, K. O. Maternal, fetal and placental regulation of placental iron trafficking. Placenta 125, 47–53, doi:10.1016/j.placenta.2021.12.018 (2022).

55 Cowell, W., Deyssenroth, M., Chen, J. & Wright, R. J. Maternal stress in relation to sex-specific expression of placental genes involved in nutrient transport, oxygen tension, immune response, and the glucocorticoid barrier. Placenta 96, 19–26, doi:10.1016/j.placenta.2020.05.004 (2020).

56 Litzky, J. F., Deyssenroth, M. A., Everson, T. M., Lester, B. M., Lambertini, L., Chen, J. et al. Prenatal exposure to maternal depression and anxiety on imprinted gene expression in placenta and infant neurodevelopment and growth. Pediatr Res 83, 1075–1083, doi:10.1038/pr.2018.27 (2018).

57 Olivier, J. D., Akerud, H., Skalkidou, A., Kaihola, H. & Sundstrom-Poromaa, I. The effects of antenatal depression and antidepressant treatment on placental gene expression. Front Cell Neurosci 8, 465, doi:10.3389/fncel.2014.00465 (2014).

58 Dong, Y., Weng, J., Zhu, Y., Sun, D., He, W., Chen, Q. et al. Transcriptomic profiling of the developing brain revealed cell-type and brain-region specificity in a mouse model of prenatal stress. BMC Genomics 24, 86, doi:10.1186/s12864-023-09186-8 (2023).

59 Wang, X., Zhou, F. Y., Wu, T., Duan, C., Luo, X., Meng, Y. et al. Maternal gut microbiota mediates prenatal stress-induced fetal blood brain barrier dysfunction. Gut Microbes 18, 2631242, doi:10.1080/19490976.2026.2631242 (2026).

60 Cisse, Y. M., Montgomery, K. R., Zierden, H. C., Hill, E. M., Kane, P. J., Huang, W. et al. Maternal preconception stress produces sex-specific effects at the maternal:fetal interface to impact offspring development and phenotypic outcomesdagger. Biol Reprod 110, 339–354, doi:10.1093/biolre/ioad156 (2024).

61 Howerton, C. L., Morgan, C. P., Fischer, D. B. & Bale, T. L. O-GlcNAc transferase (OGT) as a placental biomarker of maternal stress and reprogramming of CNS gene transcription in development. Proc Natl Acad Sci U S A 110, 5169–5174, doi:10.1073/pnas.1300065110 (2013).

62 Nugent, B. M., O’Donnell, C. M., Epperson, C. N. & Bale, T. L. Placental H3K27me3 establishes female resilience to prenatal insults. Nat Commun 9, 2555, doi:10.1038/s41467-018-04992-1 (2018).

63 Behura, S. K., Kelleher, A. M. & Spencer, T. E. Evidence for functional interactions between the placenta and brain in pregnant mice. FASEB J 33, 4261–4272, doi:10.1096/fj.201802037R (2019).

64 Crampton, S. J., Collins, L. M., Toulouse, A., Nolan, Y. M. & O’Keeffe, G. W. Exposure of foetal neural progenitor cells to IL-1beta impairs their proliferation and alters their differentiation - a role for maternal inflammation? J Neurochem 120, 964–973, doi:10.1111/j.1471-4159.2011.07634.x (2012).

65 Zhang, Z., Li, N., Chen, R., Lee, T., Gao, Y., Yuan, Z. et al. Prenatal stress leads to deficits in brain development, mood related behaviors and gut microbiota in offspring. Neurobiol Stress 15, 100333, doi:10.1016/j.ynstr.2021.100333 (2021).

66 Kawamura, T., Chen, J., Takahashi, T., Ichitani, Y. & Nakahara, D. Prenatal stress suppresses cell proliferation in the early developing brain. Neuroreport 17, 1515–1518, doi:10.1097/01.wnr.0000236849.53682.6d (2006).

